# The mitochondrial genomes of 11 aquatic macroinvertebrate species from Cyprus

**DOI:** 10.1101/2020.06.26.173625

**Authors:** Jan-Niklas Macher, Katerina Drakou, Athina Papatheodoulou, Berry van der Hoorn, Marlen Vasquez

## Abstract

Aquatic macroinvertebrates are often identified based on morphology, but molecular approaches like DNA barcoding, metabarcoding and metagenomics are increasingly used for species identification. These approaches require the availability of DNA references deposited in public databases. Here we report the mitochondrial genomes of 11 aquatic macroinvertebrates species from Cyprus, a European Union island country in the Mediterranean. Only three of the molecularly identified species could be assigned to a species name, highlighting the need for taxonomic work that leads to the formal description and naming of species, and the need for further genetic work to fill the current gaps in reference databases containing aquatic macroinvertebrates.

**Graphical Abstract:** 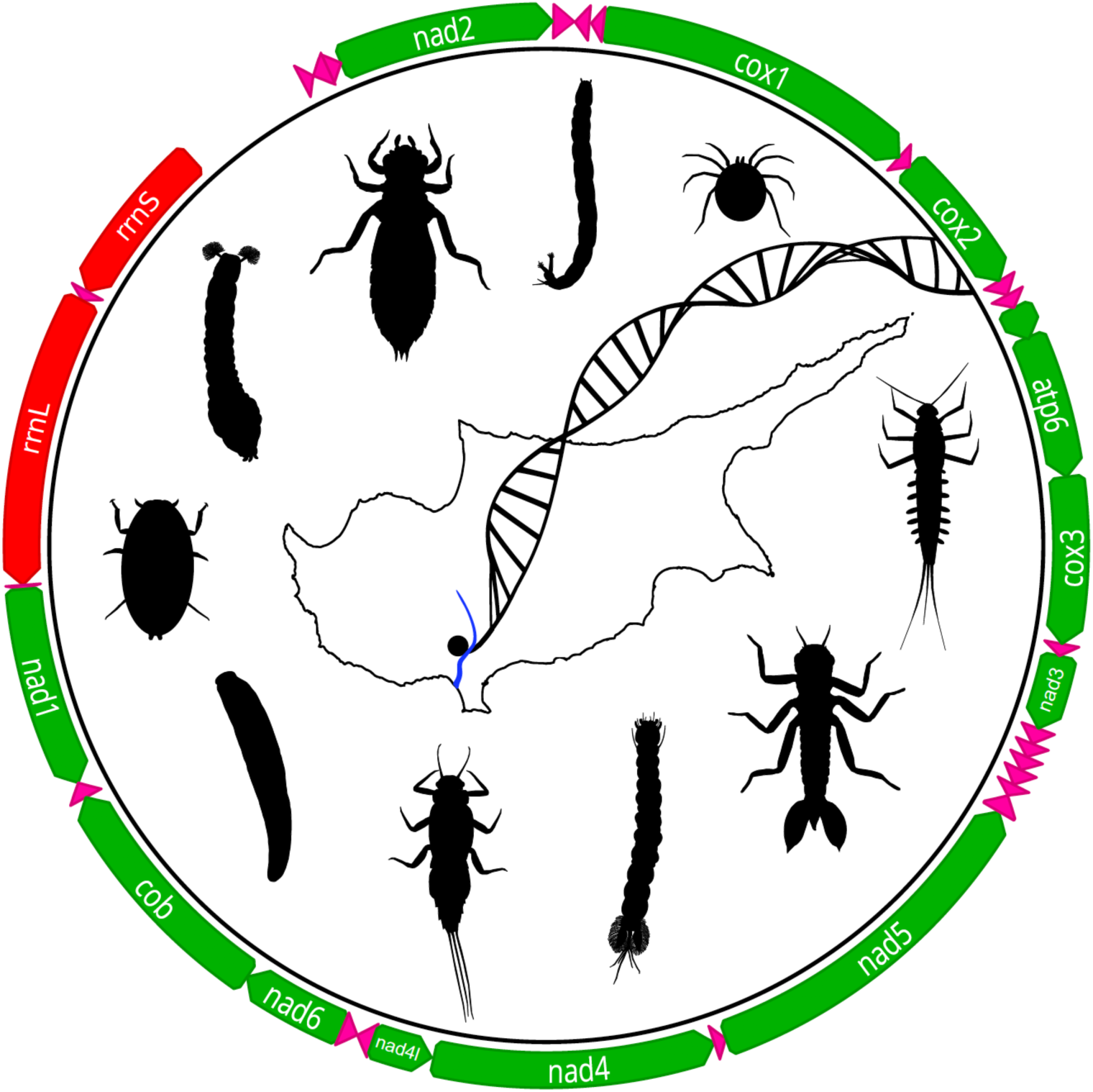

## Introduction

Aquatic macroinvertebrates are commonly used for the assessment of ecosystem quality (*1*–*3*) and in ecological studies (*4*–*6*). Taxa are often identified based on morphology, but the advent of DNA sequencing technologies has led to a paradigm shift, as techniques like DNA barcoding (*7*), metabarcoding (*8*) and metagenomic approaches (*9*–*11*) allow to identify and study large numbers of macroinvertebrate specimens, species and communities in a short amount of time and at reasonable costs. DNA based identification of macroinvertebrates relies on the comparison of obtained DNA sequence data to reference sequences deposited in databases like BOLD (*12*) or NCBI GenBank (*13*). Recent work has shown that some geographic areas and taxonomic groups are well represented in reference databases, but information on other areas and taxa is scarce, limiting the applicability of DNA based methods for studying the respective taxonomic groups and ecosystems (*14, 15*). It is therefore important to sequence aquatic macroinvertebrate species and deposit the information in public databases. Full mitochondrial genomes have the advantage of containing (usually) 13 protein coding genes and the large and small subunits of the ribosomal RNA, which can potentially help improve species identification and makes data available for phylogenetic studies.

Here we report the mitochondrial genomes of 11 aquatic macroinvertebrates species from Cyprus, a European Union island country in the Mediterranean. The aquatic macroinvertebrate fauna of Cyprus is expected to be diverse and harbour many endemic species due to the country’s location in the Mediterranean Biodiversity Hotspot (*16*). However, the fauna is vastly understudied with respect to aquatic macroinvertebrates, with less than 50 molecular barcodes deposited in the Barcode of Life database as of May 2020, and no country-specific identification keys available to date. We sequenced and assembled the mitochondrial genomes of two Baetidae (Ephemeroptera), one Caenidae (Ephemeroptera), one Chironomidae (Diptera), one Dixidae (Diptera), one Simuliidae (Diptera), one Hydrachnidae (Acari), one Gomphidae (Odonata), one Euphaeidae (Odonata), one Gyrinidae (Coleoptera) and one Erpobdellidae (Hirudinea) species. The sequencing of mitochondrial genomes of selected taxa is part of a COST DNAaqua-NET(*17*) project on Cyprus freshwater biodiversity, which tackles the challenges of describing aquatic diversity in understudied regions and making data available for future studies and biomonitoring (*18, 19*). The ongoing project will lead to an in-depth description of the aquatic macroinvertebrate fauna of Cyprus.

## Material & Methods

Specimens were collected from a perennial stream in Cyprus (coordinates: 34.769801 N, 32.911568 E, see Figure 1) on 14-08-2018 using a surber sampler (500um mesh bag). Specimens were identified on family level based on morphology by local experts. All specimens were stored in 96% ethanol and shipped to the Naturalis Biodiversity Center.

**Figure 1:**
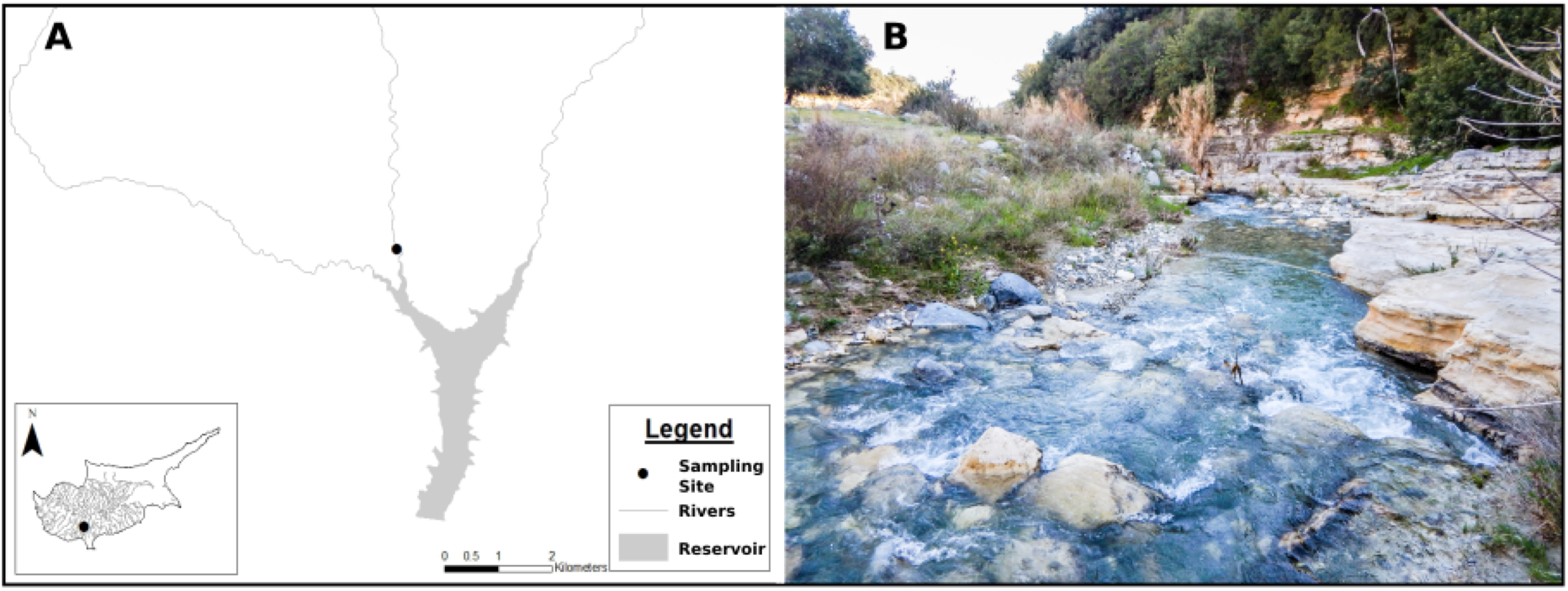
A) Location of the sampling site in Cyprus B) Photo of the sampling site

In the laboratory, specimens were carefully rinsed with lab grade water to remove external adhesions. DNA was extracted from the whole body as follows. The specimens were cut with scissors and crushed with spatulas in 2ml Eppendorf tubes. All equipment was sterile. DNA was extracted using the Macherey-Nagel (Düren, Germany) NucleoSpin tissue kit on the KingFisher (Waltham, USA) robotic platform, following the manufacturer’s protocol. A negative control containing lab grade water was processed together with the tissue samples and was checked for contamination during all following steps. Extracted DNA was stored at -20 °C overnight until further processing. Quantity and size of the extracted DNA was checked on the QIAxcel platform. 15ng of DNA per sample was used for further processing. The New England Biolabs (Ipswitch, USA) NEBNext kit and oligos were used for library preparation, following the manufacturer’s protocol and selecting for an average DNA fragment length of 300 bp. Fragment length and quantity of DNA in samples was checked on the Agilent (Santa Clara, USA) Bioanalyzer. Samples were equimolar pooled before sending for sequencing. The negative control, which did not show any signal of DNA present, was added to the final library with 10% of the total library volume. The final library was shipped to BGI (Shenzhen, China) for sequencing on the NovaSeq 6000 platform (2x 150bp).

Raw reads were quality checked using MultiQC (*20*). Fastp (*21*) was used for trimming of Illumina adapters and removal of reads shorter than 100 bp. Reads were assembled using megahit (*22*) with a maximum k-mer length of 69 and default parameters. Spades (*23*) was used for a separate assembly to assess consistency of assembly results (all commands: Supplementary material 1). Contigs were blast searched against the NCBI Genbank nucleotide database (*13*). Identified putative mitochondrial genomes were annotated using the MITOS (*24*) web server. Reading frames and annotations were manually checked and corrected using Geneious Prime (v. 2020.1), which was subsequently used to submit the annotated mitochondrial genomes to NCBI Genbank. Gene order and direction was compared to the available mitochondrial genomes of related species downloaded from NCBI Genbank. The obtained cytochrome c oxidase I (COI) genes were compared with sequences deposited in the NCBI Genbank and BOLD databases to check for availability of species-level barcodes. A match of 97% identity or higher was used as threshold for a species-level match (*7*). Occurrence data was compared to literature and to data available in the Global Biodiversity Information Facility.

## Results

Sequencing on the NovaSeq platform resulted in 43,369,218 reads (reads per sample: Supplementary table 1). The negative control contained 952 reads (0.002% of all). Illumina platforms are known to produce tag switching (*25, 26*), and a small proportion of reads is commonly found in negative controls. As the number of reads observed in our negative control was low, we did not suspect contamination. We obtained the full mitochondrial genome for all 11 macroinvertebrate species. Coverage varied from 18 (Euphaeidae_Cy2020_sp. 1, Odonata) to 449 (Caenidae_Cy2020_sp. 1, Ephemeroptera), with an average coverage of 135-fold (see supplementary table 2 for coverage and assembly length). All mitochondrial genomes contained 13 protein coding genes, 22 tRNAs and the large and small subunit of the ribosomal RNA (see supplementary material 2 for mitochondrial genomes. Fully annotated genomes will be available in NCBI GenBank upon publication). Megahit and Spades assemblies resulted in identical mitochondrial contigs, with the expected exception being length variations in the highly repetitive and AT rich control regions, which are difficult to assemble with short reads. Gene order and direction of all mitochondrial genomes was identical to that of related taxa available in Genbank.

**Table 1:**
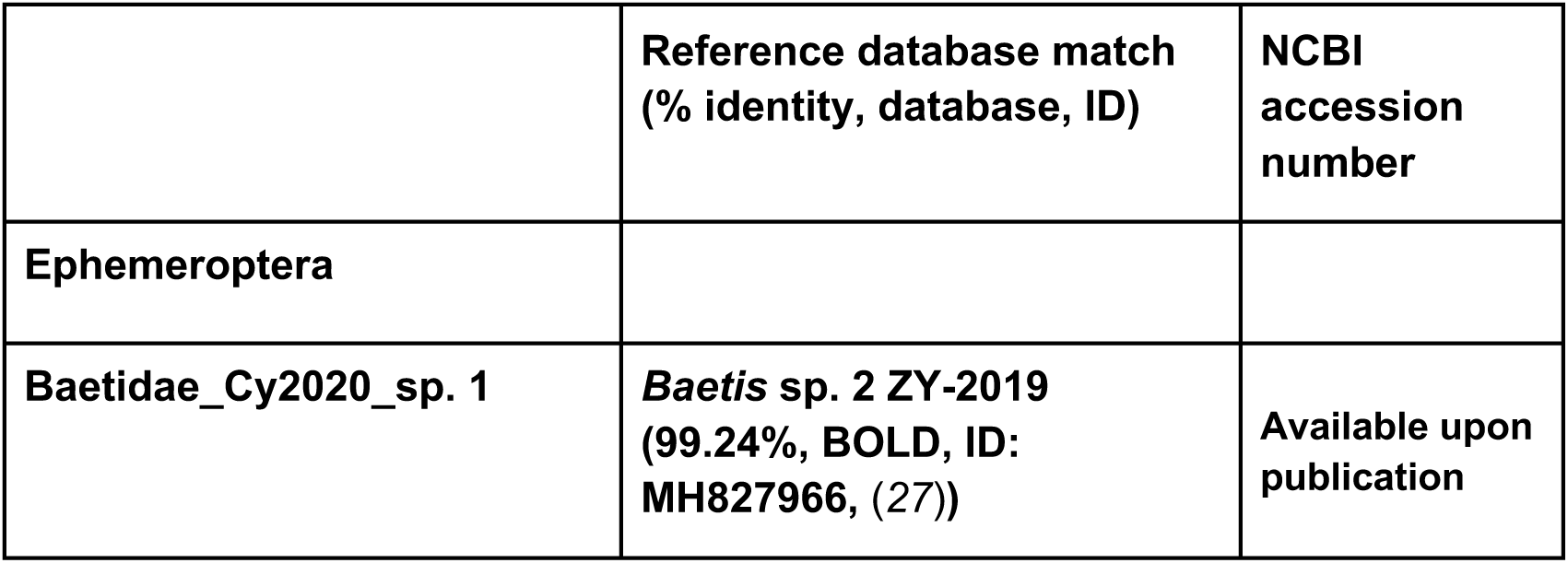
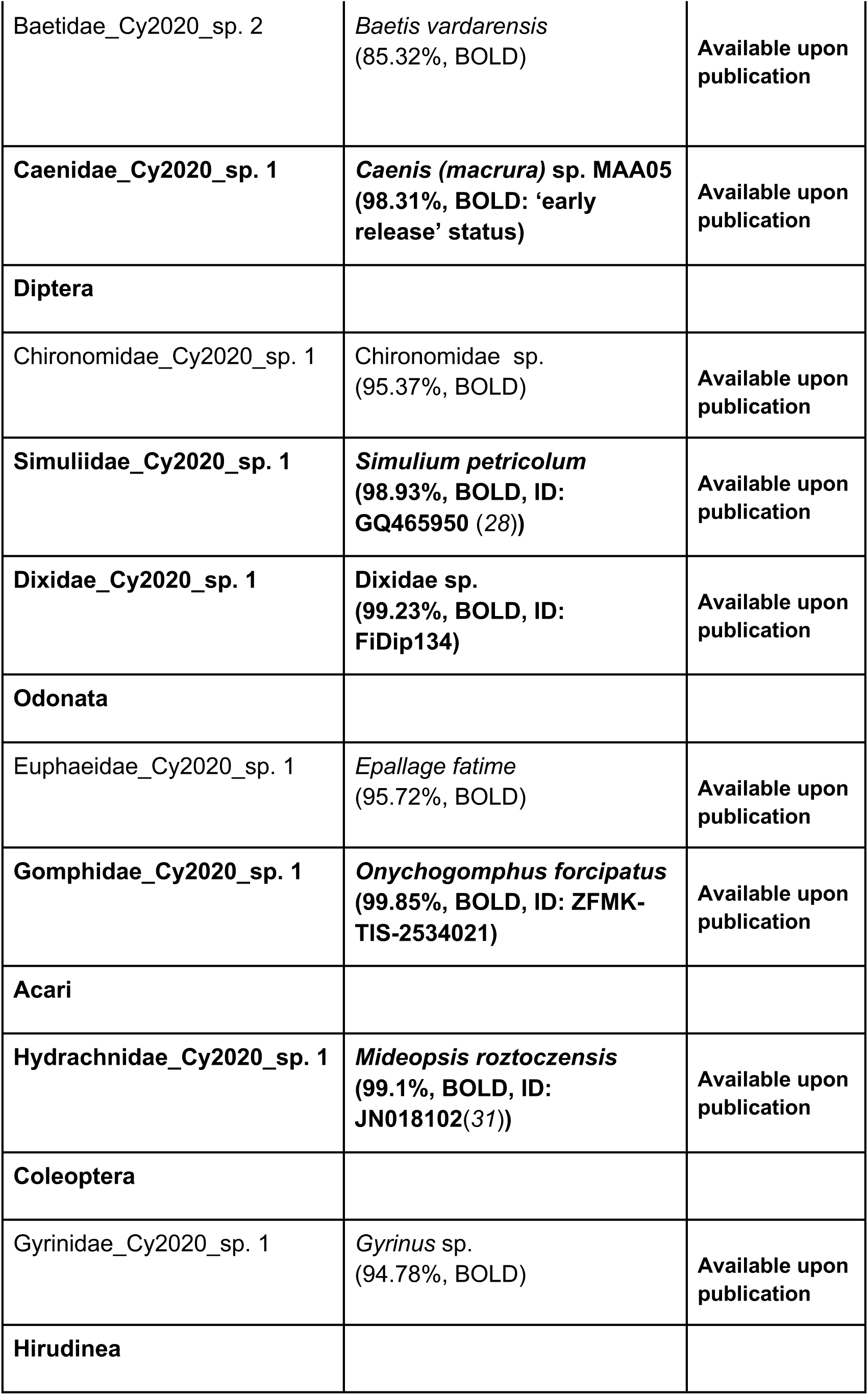
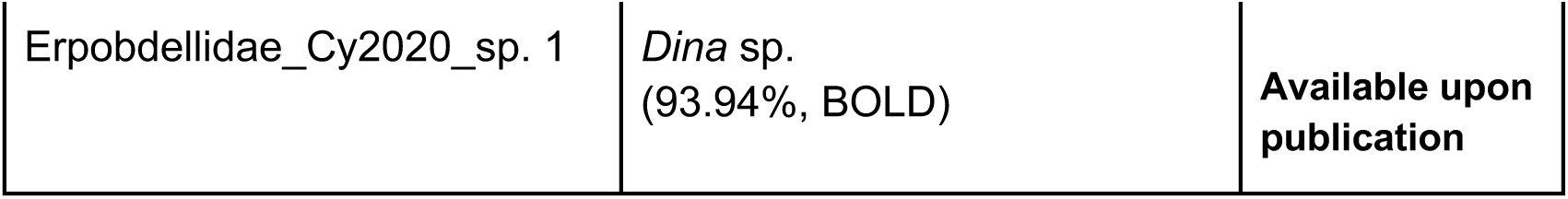
Blast results and NCBI accession numbers for the 11 assembled mitochondrial genomes. Identities >97% were recorded as species level match (highlighted in bold).

|  | Reference database match<br>(% identity, database, ID) | NCBI<br>accession<br>number |
| --- | --- | --- |
| **Ephemeroptera** |  |  |
| **Baetidae_Cy2020_sp. 1** | **Baetis sp. 2 ZY-2019**<br>**(99.24%, BOLD, ID:**<br>**MH827966, (27))** | **Available upon**<br>**publication** |
| Baetidae_Cy2020_sp. 2 | *Baetis vardarensis*<br>(85.32%, BOLD) | Available upon publication |
| Caenidae_Cy2020_sp. 1 | *Caenis (macrura)* sp. MAA05<br>(98.31%, BOLD: 'early release' status) | Available upon publication |
| **Diptera** |  |  |
| Chironomidae_Cy2020_sp. 1 | Chironomidae sp.<br>(95.37%, BOLD) | Available upon publication |
| Simuliidae_Cy2020_sp. 1 | *Simulium petricolum*<br>(98.93%, BOLD, ID: GQ465950 (28)) | Available upon publication |
| Dixidae_Cy2020_sp. 1 | Dixidae sp.<br>(99.23%, BOLD, ID: FiDip134) | Available upon publication |
| **Odonata** |  |  |
| Euphaeidae_Cy2020_sp. 1 | *Epallage fatime*<br>(95.72%, BOLD) | Available upon publication |
| Gomphidae_Cy2020_sp. 1 | *Onychogomphus forcipatus*<br>(99.85%, BOLD, ID: ZFMK-TIS-2534021) | Available upon publication |
| **Acari** |  |  |
| Hydrachnidae_Cy2020_sp. 1 | *Mideopsis roztoczensis*<br>(99.1%, BOLD, ID: JN018102(31)) | Available upon publication |
| **Coleoptera** |  |  |
| Gyrinidae_Cy2020_sp. 1 | *Gyrinus* sp.<br>(94.78%, BOLD) | Available upon publication |
| **Hirudinea** |  |  |
| Erpobdellidae_Cy2020_sp. 1 | *Dina* sp.<br>(93.94%, BOLD) | Available upon<br>publication |

Of the 11 species, six could be molecularly identified on species level using the 97% identity threshold. However, only three of these six species could be assigned to a full binomial name, while the other three were assigned to references with preliminary names, pending formal description or taxonomic identification (see table 1). ‘Baetis sp. 2 ZY-2019’ was previously identified in Israel (*27*), which is close to Cyprus. ‘Caenis (macrura) sp. MAA05’ was reported from streams in Northern Iraq (BOLD ID: BMIKU-0029). The dipteran *Simulium petricolum* was previously reported from areas around the Mediterranean and from Britain (*28*). The Dixidae species with BOLD ID FiDip134 was previously found in Norway, but only two reference sequences from the same region are available and a lack of a formal species name does not allow assessing whether this species has previously been found in the mediterranean area. The dragonfly *Onychogomphus forcipatus* is known from a wide range across Europe, including the mediterraenan areas (occurrence data from GBIF: (*29*)), and the water mite *Mideopsis roztoczensis* is known from areas in central Europe as well as from the areas around the Mediterranean (GBIF data: (*30*)). These preliminary results indicate that the aquatic macroinvertebrate fauna of Cyprus consists of both European and Asian species.

Our results highlight the need for taxonomic work that leads to the formal description and naming of species, and the need for further genetic work to fill the current gaps in reference databases containing aquatic macroinvertebrates (*14, 15*). This is especially important for yet understudied regions like Cyprus, for which no specific identification keys for macroinvertebrates exist. We show that shotgun sequencing can be used for rapid recovery of mitochondrial genomes and can help with filling the gaps in reference databases. The ongoing COST DNAaqua-NET (*17*) project on Cyprus freshwater biodiversity will combine taxonomic and molecular approaches and will lead to an in-depth description of the aquatic macroinvertebrate fauna of Cyprus.

## Acknowledgements

We thank Elza Duijm, Frank Stokvis, Marcel Eurlings and Roland Butôt for support in the lab.

This article is based upon work from COST Action DNAqua-Net (CA15219), supported by the COST (European Cooperation in Science and Technology) program.

## Data availability

All raw data has been deposited in the NCBI Short Read Archive (SRA). Accession numbers: Available upon publication. All mitogenomes have been deposited in NCBI GenBank. Accession numbers: Available upon publication. All DNA extracts are stored in the Naturalis Biodiversity Center collection.

