## Supplementary tables 1 & 2 for "The mitochondrial genomes of 11 aquatic macroinvertebrate species from Cyprus"

| Supplementary table 1: No. of raw reads per sample, and reads retained after quality filtering | | | |
|---|---|---|---|
| **Species** | **Raw reads** | **Read no. after qc** | **% of reads retained after qc** |
| **Baetidae_Cy2020_sp. 1** | 2565416 | 2481508 | 96.7292634 |
| **Baetidae_Cy2020_sp. 2** | 4552074 | 4411408 | 96.90984813 |
| **Chironomidae_Cy2020_sp. 1** | 2598816 | 2538078 | 97.66285878 |
| **Caenidae_Cy2020_sp. 1** | 2448332 | 2333082 | 95.29271357 |
| **Dixidae_Cy2020_sp. 1** | 3664936 | 3569352 | 97.39193263 |
| **Gomphidae_Cy2020_sp. 1** | 3935716 | 3478546 | 88.3840704 |
| **Gyrinidae_Cy2020_sp. 1** | 7297418 | 6901072 | 94.56868169 |
| **Hydrachnidae_Cy2020_sp. 1** | 3203460 | 3130448 | 97.72083934 |
| **Erpobdellidae_Cy2020_sp. 1** | 2705890 | 2456086 | 90.76813913 |
| **Euphaeidae_Cy2020_sp. 1** | 6628714 | 4764394 | 71.87508769 |
| **Simuliidae_Cy2020_sp. 1** | 3768446 | 3701984 | 98.23635525 |
| **Negative control** | 952 | | |

| Supplementary table 2: Mean coverage and length of mitochondrial genomes | | |
|---|---|---|
| **Species** | **Mean coverage** | **Length (bp)** |
| **Euphaeidae_Cy2020_sp. 1** | 18 | 15251 |
| **Gomphidae_Cy2020_sp. 1** | 30 | 14917 |
| **Erpobdellidae_Cy2020_sp. 1** | 50 | 14746 |
| **Chironomidae_Cy2020_sp. 1** | 61 | 17345 |
| **Hydrachnidae_Cy2020_sp. 1** | 72 | 13989 |
| **Simuliidae_Cy2020_sp. 1** | 105 | 15888 |
| **Dixidae_Cy2020_sp. 1** | 115 | 15472 |
| **Gyrinidae_Cy2020_sp. 1** | 143 | 16178 |
| **Baetidae_Cy2020_sp. 1** | 171 | 15391 |
| **Baetidae_Cy2020_sp. 2** | 267 | 15,086 |
| **Caenidae_Cy2020_sp. 1** | 449 | 15658 |
