## Supplementary material 1 for "The mitochondrial genomes of 11 aquatic macroinvertebrate species from Cyprus"

**###Assembly of mitogenomes**
**##Commands shown for one sample**

##fastp adapter trim and removal of sequences <100bp

```
fastp -i Simuliidae_Cy2020_sp1_1.fq.gz -o  Simuliidae_Cy2020_sp1_1_trimmed.fq.gz    \
 -I Simuliidae_Cy2020_sp1_2.fq.gz  -O Simuliidae_Cy2020_sp1_2_trimmed.fq.gz     \
 --detect_adapter_for_pe --length_required 100 -x
```

##megahit assembly

```
megahit -1 Simuliidae_Cy2020_sp1_1_trimmed.fq.gz  -2
Simuliidae_Cy2020_sp1_2_trimmed.fq.gz  -t 20 --min-contig-len 1000 -o
Simuliidae_Cy2020_sp1 —k-list 21,29,39,49,59,69
```

##Spades assembly

```
spades.py -o ~/Spades/output -1 Simuliidae_Cy2020_sp1_1_trimmed.fq.gz -2
Simuliidae_Cy2020_sp1_2_trimmed.fq.gz -t 20
```
